# High-quality genome-scale metabolic model of *Aurantiochytrium* sp. T66

**DOI:** 10.1101/2020.11.06.371070

**Authors:** Vetle Simensen, André Voigt, Eivind Almaas

## Abstract

The long-chain, *ω*-3 polyunsaturated fatty acids (PUFAs) eicosapentaenoic acid (EPA) and docosahexaenoic acid (DHA) are essential for humans and animals, including marine fish species. Presently, the primary source of these PUFAs is fish oils. As the global production of fish oils appears to be reaching its limits, alternative sources of high-quality *ω*-3 PUFAs is paramount to support the growing aquaculture industry. Thraustochytrids are a group of heterotrophic protists able to synthesize and accrue large amounts of essential *ω*-3 PUFAs, including EPA and DHA. Thus, the thraustochytrids are prime candidates to solve the increasing demand for *ω*-3 PUFAs using microbial cell factories. However, a systems-level understanding of their metabolic shift from cellular growth into lipid accumulation is, to a large extent, unclear. Here, we reconstructed a high-quality genome-scale metabolic model of the thraustochytrid *Aurantiochytrium* sp. T66 termed iVS1191. Through iterative rounds of model refinement and extensive manual curation, we significantly enhanced the metabolic scope and coverage of the reconstruction from that of previously published models, making considerable improvements with stoichiometric consistency, metabolic connectivity, and model annotations. We show that iVS1191 is highly consistent with experimental growth data, reproducing *in vivo* growth phenotypes as well as specific growth rates on minimal carbon media. The availability of iVS1191 provides a solid framework for further developing our understanding of T66’s metabolic properties, as well as exploring metabolic engineering and process-optimization strategies *in silico* for increased *ω*-3 PUFA production.

## Introduction

An increasing amount of evidence have been gathered during the last decades of the health benefits from marine seafood consumption (Hosomi, Yoshida, & Fukunaga, 2012). The high content of *ω*-3 polyunsaturated fatty acids (PUFAs), such as docosahexaenoic acid (DHA), is considered one of the major contributors to this effect (Hosomi et al., 2012; Swanson, Block, & Mousa, 2012). These fatty acids are necessary for proper development of the fetal brain and retina (Coletta, Bell, & Roman, 2010; Lauritzen et al., 2016), and have been shown to possess anti-inflammatory properties associated with reduced risks of cardiovascular disease (Bloomer, Larson, Fisher-Wellman, Galpin, & Schilling, 2009; Bouwens et al., 2009). Although rich in *ω*-3 fatty acids, oily fish such as salmon do not synthesize these *de novo*, but rather acquire them from feeding on lipid-rich plankton and microalgae (Hosomi et al., 2012). The rising global aquaculture industry, particularly the marine pisciculture industry, has consequently led to an increasing need for novel sources of these fatty acids (Food and Agriculture Organization of the United Nations, 2018), the current source of which is mainly fish oil. Due to their trophic level in the food chain, these PUFA-containing lipids also contain elevated levels of toxic contaminants, such as polychlorinated biphenyls and heavy metals (El-Moselhy, Othman, Abd El-Azem, & El-Metwally, 2014; Rawn et al., 2006). A need for alternative, clean, high-quality sources of *ω*-3 PUFAs is therefore of utmost importance.

Cultivation and strain engineering of lipid-producing microorganisms is regarded as a promising strategy to cater to this demand. These oleaginous organisms are able to accumulate substantial amounts of lipids and has proven to be a cost-effective alternative to agricultural oil production (Ratledge, 2004). Despite the fact that a large variety of phylogenetically diverse microorganisms are able to produce and store lipids, only a limited number of oleaginous species produce lipids rich in *ω*-3 PUFAs (Gupta, Barrow, & Puri, 2012). One of these is a taxonomically ambiguous group of heterokonts called thraustochytrids. The thraustochytrids are heterotrophic protists found globally in marine and estuarine environments (Leyland, Leu, & Boussiba, 2017). They accrue large quantities of *ω*-3-rich triacylglycerols (TAGs) stored as intracellular lipid droplets that may constitute 50-80% of the total cell mass (Huang, Lu, & Chu, 2012; Leyland et al., 2017; Li et al., 2015). Although the fatty acid composition of these TAGs vary between different thraustochytrid species, they predominately contain palmitic acid and DHA (Morabito et al., 2019).

In most PUFA-producing microorganisms, the fatty acids are produced by elongating and desaturating the end products of the type I fatty acid synthase (FAS) enzymatic machinery (Beld, Lee, & Burkart, 2015). However, thraustochytrids predominately synthesize their PUFAs using a competing polyketide synthase-like (PKS) system thought to have originated from marine prokaryotes by means of lateral gene transfer (Metz et al., 2001). Here, the acyl intermediates may retain their unsaturated bonds during the biosynthetic process, thus lowering the molar requirement for reducing power compared to the conventional elongation/desaturation scheme (Ratledge, 2004). This capability to accumulate large amounts of *ω*-3 PUFAs, as well as the decreased biosynthetic requirement for reducing power, has made thraustochytrids particularly promising candidates as efficient lipid-producing cell factories (Ratledge, 2004).

To elucidate the biological mechanisms underpinning the fatty acid biosynthesis and lipid accumulation in thraustochytrids, knowledge of their global metabolic organization and functionality is vital. Metabolic modeling is a pivotal methodology to simulate and predict metabolic phenotypes *in silico*, thereby generating hypotheses and guide experimental efforts (O’Brien, Monk, & Palsson, 2015). Specifically, genome-scale metabolic models (GSMs) have become one of the main approaches to model metabolism within the field of systems biology (Gu, Kim, Kim, Kim, & Lee, 2019). Whereas some modeling approaches require extensive parametrization, constraint-based modeling using GSMs in their most fundamental form merely calls for an annotated genome sequence (Thiele & Palsson, 2010). GSMs contain the genotype-phenotype relationships for genes and biochemical reactions, enabling the computational simulations of metabolic flux distributions under various environmental and genetic perturbations (O’Brien et al., 2015).

The process of GSM reconstruction is a labour-intensive endeavor that is based on the prediction of putative metabolic functions of protein-encoding sequences using the aforementioned genome annotation (Mendoza, Olivier, Molenaar, & Teusink, 2019). Once collected, this repository of biochemical reactions and metabolites are converted into the mathematical framework of a stoichiometric matrix, enabling the simulation of metabolic phenotypes using flux balance analysis (FBA) and related approaches (Bordbar, Monk, King, & Palsson, 2014). Although consecutive in order, these stages are organized in an iterative manner in which additional model curation is performed to ensure that the predictions progressively match the expected biochemical phenotype (Thiele & Palsson, 2010). While manual curation is an integral part of achieving a high-quality GSM (Mendoza et al., 2019), many of the steps are included in fully automated software producing ready-to-use metabolic reconstructions (Agren et al., 2013; Dias, Rocha, Ferreira, & Rocha, 2015; Henry et al., 2010; Karp et al., 2016; Machado, Andrejev, Tramontano, & Patil, 2018; Pitkänen et al., 2014).

Herein, we present iVS1191, the reconstruction of a high-quality GSM of the thraustochytrid *Aurantiochytrium* sp. strain T66 (Anita N Jakobsen, Aasen, & Strøm, 2007). By comprehensively refining and curating the reconstruction, we significantly improved the metabolic scope and coverage from that of a template model. We further utilized experimental growth data, proving the model able to accurately reproduce *in vivo* metabolic growth phenotypes. Currently, iVS1191 represents the most comprehensive knowledge-base of any thraustochytrid, and we strongly believe that the reconstructed model has the potential to play a key role in understanding the oleaginous metabolism of thraustochytrids for future biotechnological applications.

## Materials and Methods

### Model simulations

We use flux balance analysis (FBA) to predict cellular phenotypes of the GSM. The general FBA problem can be stated as the following linear program:

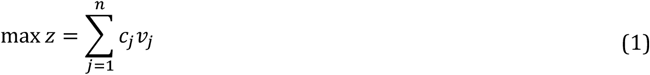

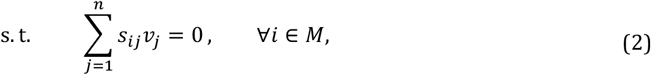

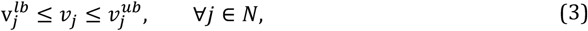

where *C_j_* is the objective function coefficient of reaction flux *V_j_* defining the objective function *z* as a linear combination of the reaction fluxes. Eq. (2) represents the homogeneous system of linear equations, while Eq. (3) specifies the upper (*v_j_^ub^*) and lower (*v_j_^lb^*) bounds on the flux variables (Orth, Thiele, & Palsson, 2010). Unless otherwise stated, all FBA simulations took the form as seen in Eq. (1)-(3) where the flux of the biomass pseudoreaction was selected as the objective. Simulations were performed on a Macbook Pro 2018 with Matlab 2019b (MATLAB, 2019) using the The COnstraint-Based Reconstruction and Analysis (COBRA) toolbox (Heirendt et al., 2019) and the proprietary solver Gurobi version 8.1.1 (Gurobi Optimization, 2020).

### Initial draft reconstruction

We used the already published GSM iCY1170_DHA (C Ye et al., 2015) of the closely related thraustochytrid *Schizochytrium limacinum* SR21 as a starting template. As our next step, we performed a reciprocal protein Basic Local Alignment Search Tool (BLASTp) (Altschul, Gish, Miller, Myers, & Lipman, 1990) analysis using the RAVEN Toolbox (Agren et al., 2013) with the predicted protein sequences of the template and target organisms as input. The protein sequences of *S. limacinum* were obtained from the Joint Genome Institute (JGI) Database (Institute, 2017). The resulting sequence similarities were subsequently used to infer putative homologs by bidirectional best hits using the following parameters: alignment length of at least 200, an E-value < 10^-30^, and a minimum identity of 40%. We also performed additional identification of candidate homologs of proteins that were unable to satisfy these similarity criteria due to short sequence lengths. All reactions from the template model associated with the identified set of putative homologs were extracted to form the initial draft reconstruction. Additionally, reactions annotated as spontaneous in the template model were also included in the draft model.

The semi-automatic gap-filling approach was carried out as follows: For every biomass precursor incapable of being produced, we studied the anabolic pathways responsible for their production in the Kyoto Encyclopedia of Genes and Genomes (KEGG) (Kanehisa & Goto, 2000) and MetaCyc (Kothari et al., 2013) databases in order to identify putative gap-filling candidates. We also utilized the software ModelExplorer (Martyushenko & Almaas, 2019) for visual and algorithmic inspection of the metabolic network to identify metabolic gaps and inconsistencies. Finally, we filled the remaining gaps by adding a minimal subset of all unincorporated template model reactions necessary for the model to generate biomass.

### Draft reconstruction improvement based on KEGG

We reconstructed a secondary draft model using the KEGG functionality of the RAVEN Toolbox (Agren et al., 2013). This method make use of profile hidden-Markov models (HMM) pre-trained on KEGG Orthologies (KO) in the KEGG database and attempts to assign these identifiers to significantly homologous genes from organism of interest. The associated KO reactions are collected and used to generate a draft reconstruction. We obtained the secondary draft model by querying the predicted protein sequences of T66 against a pre-trained profile HMM (Eukaryota, Identity: 100%) from Ref. (ToolBox, 2017).

### Manual curation of reconstructed network

We performed a manual gene re-annotation of all metabolic genes present in the two draft reconstructions by individually inspecting and evaluating their presumptive metabolic functionality. We further verified these by BLASTp searches against the KEGG (Kanehisa & Goto, 2000) and UniProtKB/Swiss-Prot (Apweiler et al., 2004) databases. Associated reactions were collected and compared to those present in the draft reconstruction. These reactions were either directly found in the databases or, when available, from appropriate genome-scale reconstructions in the BiGG database (King et al., 2015). Any discrepancies were resolved through the addition or removal of reactions and/or associated genes.

Subcellular predictions were made for all putative proteins using HECTAR (Gschloessl, Guermeur, & Cock, 2008) and DeepLoc (Almagro Armenteros, Sonderby, Sonderby, Nielsen, & Winther, 2017). In cases of differing predictions, HECTAR took precedence over DeepLoc as it is specifically tailored to predicting the subcellular targeting of proteins from the eukaryotic clade of heterokonts (Gschloessl et al., 2008; Leyland et al., 2017). We also performed qualitative evaluations in cases of conflicting predictions or when subcellular localization could be established from high-quality biochemical information. When subcellular localization of existing genes in the model was changed, appropriate transport reactions were added if subsections of the associated metabolic pathways now appeared in different compartments. We either found candidate transporter proteins directly from the genome annotation or by BLAST searches against the aforementioned databases, as well as in the Transporter Classification Database (TCDB) (Saier M. H. et al., 2016).

### Detection and removal of energy-generating cycles (EGCs)

To identify the presence of EGCs, we added a set of energy dissipation reactions ξ and tested whether these were able to carry a non-zero flux in a closed model (a listing is available in ‘SuppInfo Dissipation reactions’). Each dissipation reaction *e_l_* ∈ ξ was iteratively added to the reconstruction and subsequently selected as the objective function to be maximized in a closed model (i.e. all exchange reactions constrained to zero). This approached is described by the following linear program

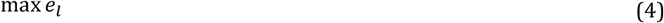

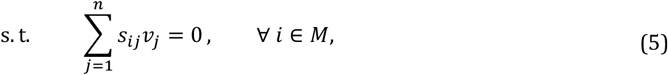

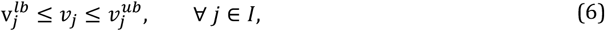

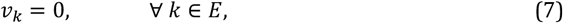

where *M* is the set of metabolites, and *I* and *E* are the sets of internal and exchange reactions, respectively. In the case of a non-zero objective value, a reaction-deletion strategy was subsequently proposed in which a minimal subset of reactions to be deleted from these flux-carrying reactions was identified. A similar test was performed with only the uptake of the carbon source constrained to zero to account for alternative EGCs in which extracellular protons are taken up and used to drive the charging of energy carriers.

### Biomass reformulation

The biomass composition was adapted from the neutral lipid-free biomass (Shene et al., 2020). Splitting the biomass into preceding precursor reactions, we formulated unique biomass reactions of the following categories: proteins, carbohydrates, DNA, RNA, unsaponifiable matter, polar lipids, and free fatty acids. The relative molar content of each amino acid in proteins was assumed to be the same as that of *Schizochytrium limacinum* SR21 (C Ye et al., 2015). We also swapped each amino acid for their corresponding aminoacyl-tRNA and added additional energy carriers required for the process of translation. The total carbohydrate content was obtained from previous work (Shene et al., 2020), while the weight fractions was assumed to be the same as that of *S. limacinum* (C Ye et al., 2015). To estimate the energy requirements for the polymerization of these monomeric carbohydrates, we employed their corresponding nucleotide sugars. As with the amino acids, we assumed that the molar abundance of each (deoxy) nucleotide in DNA and RNA was similar to that of *S. limacinum* (C Ye et al., 2015). We also substituted the monomeric precursors with corresponding (deoxy)nucleoside triphosphates, simulating the energetic cost of replication and transcription, respectively. We used previously determined levels of unsaponifiable matter, polar lipids, and free fatty acids (Shene et al., 2020), while updating the fatty acyl distribution to that of T66 cells sampled during exponential growth. Lacking detailed experimental data on the quantitative levels of specific lipids of particular chain configurations, we further assumed that all lipid classes would have the same fractional acyl chain composition. We stoichiometrically weighted the components of each of the precursor reactions using appropriate correction factors and further scaled the precursors in the final biomass reaction using experimentally determined weight fractions (i.e. g component gDW^-1^) (Shene et al., 2020).

### Phenotypic growth predictions

In order to compare *in vivo* and model-predicted *in silico* growth rates, we performed two growth experiments separately using glucose and glycerol as the carbon sources, together with ammonium as a nitrogen source in both experiments. Cells were cultivated in 100 mL medium in 500 mL shaking flasks at 28°C and 250 rpm. Four samples of 10 mL were extracted during the exponential growth phase and subsequently centrifuged at 3,200g for 10 minutes. The supernatant was frozen to quantify the amount of the carbon source and ammonium. The pellet was washed once in 0.9% NaCl, and the resulting supernatant was carefully removed before drying at 105°C for 16-20 hours to enable dry weight quantification. Glucose and glycerol were quantified by high-performance liquid chromatography (HPLC). Samples were centrifuged and filtered through 0.2 *μm* syringe filters before analyzed using an Aminex HPX-87-H column (BioRad Laboratories) at 45°C, and refractive index detection (RID-6A, Shimadzu). Five mmol H_2_SO_4_ was used as mobile phase at 0.6 mL/min. Ammonium quantities were determined using an enzymatic kit according to the manufacturer’s instructions (Megazyme K-AMIAR). Quantified levels of the carbon source and ammonium were subsequently used to estimate the specific substrate uptake rates.

## Results

### Initial draft reconstruction

We initialized the reconstruction of the metabolic model by performing genome-wide bidirectional BLAST searches for orthologs between the template model proteins of *Schizochytrium limacinum* SR21 and the predicted protein sequences of T66. In total, 995 candidate homologs were identified, which along with 1,344 associated reactions and 1,515 metabolites formed the basis of the draft reconstruction. Additionally, 87 putative homologs were found using reciprocal protein-BLAST searches with less stringent alignment length cutoffs. We subsequently used the remaining set of template genes with no significant hits in tBLASTn searches against the genomic nucleotide sequence of T66 to circumvent any insufficient genome coverage of the predicted protein sequences. A total of 30 additional candidate orthologs were identified, which along with the previous set of 87 proteins added 88 new metabolic reactions to the draft reconstruction. We also added all the unincorporated non-gene-associated transport and exchange reactions of the template model to the draft reconstruction. This was done to prevent their exclusion from impeding the ability of the model to generate necessary biomass precursors. At a later stage, we evaluated each of these reactions individually and removed those that were unnecessary for a functional model.

To ensure that the model would be able to predict specific growth rates, we initiated a semi-automatic approach to identify and fill in the metabolic gaps necessary for biomass production. Due to insufficient knowledge of the particular biomass composition of T66, the biomass reaction of the template model was employed during these simulations. Although the primary goal was to enable the synthesis of biomass, we also introduced additional reactions to more accurately represent the metabolic network of T66. In total, we added 131 reactions during this procedure of which five were non-spontaneous gap-filling reactions with no identifiable candidate genes. Two additional gap-filling reactions were found as a minimal subset of reactions from iCY1170_DHA necessary for the synthesis of the biomass precursor L-galactose. These reactions were GDP mannose phosphorylase (EC:2.7.7.22) and GDP-L-galactose phosphorylase (EC:2.7.7.69). Interestingly, both of these gap-filling reactions had no associated genes in the template model, suggesting that their responsible enzymes might be elusive in thraustochytrids in general.

### Draft reconstruction from the KEGG database

To increase the predictive capabilities of the newly reconstructed model, we initiated comprehensive investigations of additional metabolic functionalities. To facilitate this process and aid in the identification of novel reactions and genes, we generated a secondary draft model from the KEGG database (Kanehisa & Goto, 2000) using the RAVEN Toolbox (Agren et al., 2013). The contents of this model, along with biochemical information available within the genome annotation were then used to identify reactions and genes that should be added to the draft reconstruction. We also identified candidate metabolic reactions through the inspection of metabolic dead-ends and associated blocked reactions of the model. Candidate genes for missing reactions were thereby identified and suitable metabolic reactions were collected, forming a repository of novel metabolic capability to be added to the draft reconstruction.

The secondary draft model harboured 1,455 reactions, 1,556 metabolites, and 1,090 associated genes. Of these genes, 340 were unique to the KEGG model, while 404 genes where unique to the initial draft model. This highlights quite a substantial difference in genetic coverage between the two models and implies that the metabolic potential of both reconstructions might be inadequate. A similar discrepancy was found when studying the subsystem distributions of the model reactions in Figure 1. Although the number of reactions associated with each subsystem appear to be somewhat comparable, the sets of unique reactions make up a significant amount of all model reactions. In total, the initial draft reconstruction and the KEGG model each include 54.6% and 55.1% reactions that are not present in the other model. In the case of the draft reconstruction, these reactions were predominately associated with the transport and exchange subsystems. While the KEGG database contains a comprehensive repository of high-quality biochemical information, it is not specifically tailored to assist in the reconstruction of GSMs. This is partly reflected in the absence of implemented transport reactions, which plays a key role in the genesis of genome-scale metabolic networks, particularly for the compartmentalized metabolism of eukaryotes. Associated with this is also information on the subcellular localization of proteins, which altogether is lacking in the model generated from KEGG.

**Figure 1:**
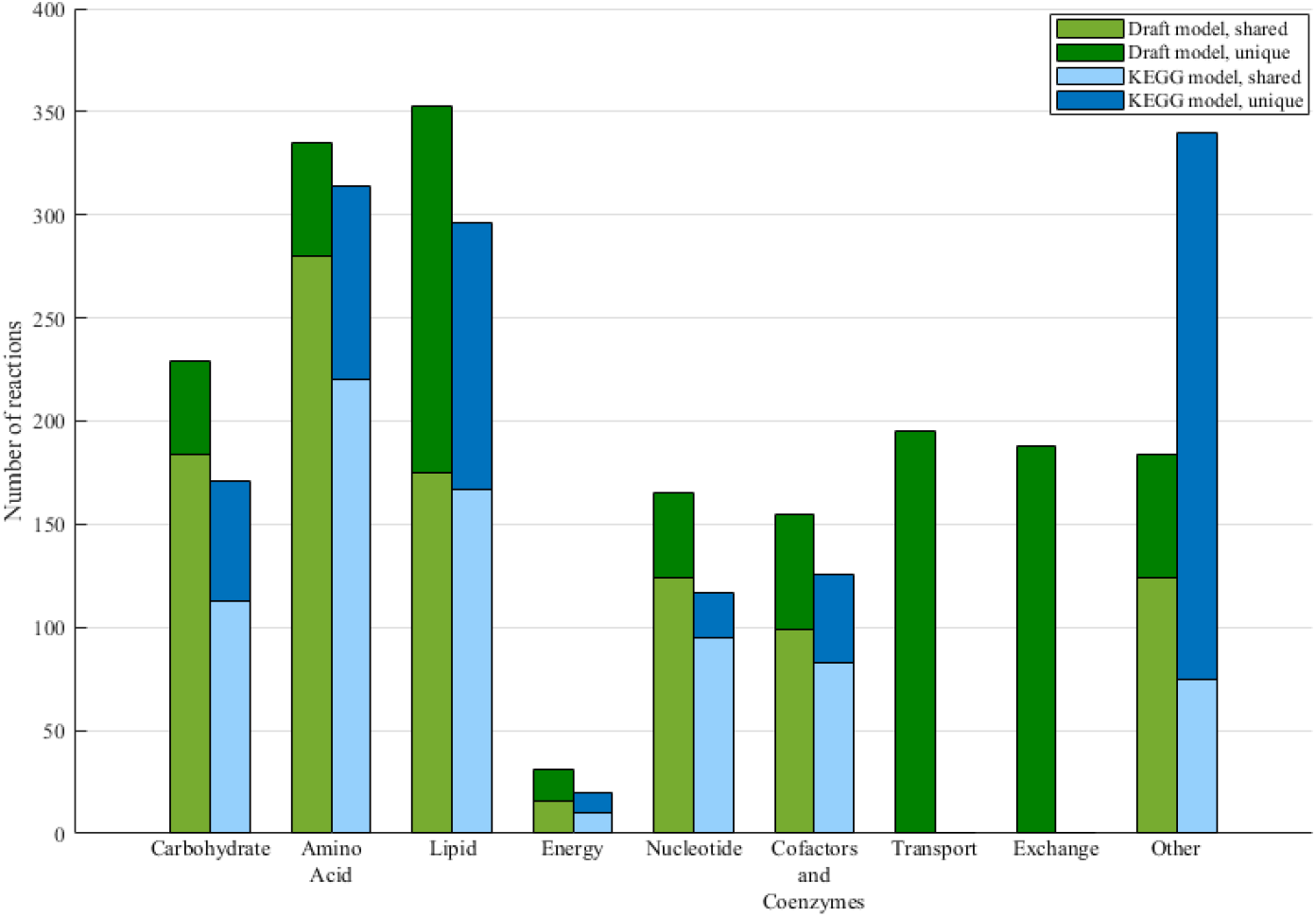
Subsystem distributions of the shared and unique reactions in the initial draft reconstruction of T66 and the KEGG model. The dissimilar number of shared reactions in each subsystem is a consequence of certain reactions of the draft reconstruction occurring in multiple compartments, or that a shared reaction are annotated to different subsystems in the two models.

### Manual curation

Motivated by the large metabolic disparity of the two reconstructions, we began a comprehensive model consolidation process. Here, the aim was to merge the two reconstructions while simultaneously performing manual curation on all the included model components. This entailed extensive gene re-annotation to enhance the accuracy of the gene-reaction associations, as well as assuring quality-control of the newly identified genes and associated reactions. Although highly laborious, the manual gene re-annotation proved to be a fruitful strategy. High-quality annotations on all 1191 genes of the final model were collected, strengthening the qualitative properties of the GSM and reinforcing the confidence in the included reactions compared to that of the automatically assembled draft reconstructions. The repository contains information on the presumed metabolic capabilities of all the metabolic genes, along with predicted subcellular localizations, bibliomic references of similar metabolic enzymes, and associated model reactions (see ‘SuppInfo Gene reannotation’). When building this repository, we updated the gene rules for a substantial set of the model reactions. While, in many cases the associated reactions of a given gene were accurate according to the available biochemical information, the gene rules were frequently incorrect, often as a result of the exclusion of auxiliary subunits of both regulatory and catalytic activity. Their identification during the re-annotation allowed for more realistic gene-reaction associations, which will greatly increase the validity of future *in silico* analysis of gene essentiality, as well as the identification of realistic genetic engineering strategies.

We performed extensive validation of Enzyme Commission (EC) numbers as many of those included from the template reconstruction were either erroneous or originated from obsolete and/or outdated database entries. By using the MetaCyc (Kothari et al., 2013) database as a benchmark, we validated the metabolite formulas and charges of all model metabolites, as well as the reaction stoichiometries, reversibilities, and co-factor specificities. We further added reaction identifiers from KEGG (Kanehisa & Goto, 2000), BiGG (King et al., 2015), MetaCyc (Kothari et al., 2013), and MetaNetX (Ganter, Bernard, Moretti, Stelling, & Pagni, 2013). Similarly for metabolites, we incorporated identifiers from the aforementioned database namespaces, as well as from ModelSEED (Henry et al., 2010), International Chemical Identifier (InChI) (McNaught, 2008), Chemical Entities of Biological Interest (ChEBI) (Degtyarenko et al., 2008), and Human Metabolome Database (HMDB) (Wishart et al., 2007).

Stoichiometric models of genome-scale metabolism may contain thermodynamically infeasible cycles which are able to charge intracellular energy carriers without any nutrient consumption (Fritzemeier, Hartleb, Szappanos, Papp, & Lercher, 2017). These cycles are products of inaccurate reversibilities of the constituent reactions, which often result from a lack of thermodynamic constraints. Of particular relevance are so-called energy-generating cycles (EGCs), a subset of type II extreme pathways where the cycle drive the biochemical charging of metabolic energy carriers without any input of energy (Fritzemeier et al., 2017; Price, Famili, Beard, & Palsson, 2002). Following the iterative addition and subsequent maximization of the set of energy-dissipation reactions, it was revealed that the reconstructed model was able to charge all of the energy carriers without any input of reduced carbon. These thermodynamically infeasible cycles concertedly operated by taking up extracellular protons by means of a flux coupling between a proton-driven transport of a given metabolite and a reversible uniport reaction of the same compound. The set of reactions utilized this flow of protons to drive a continuous generation of cytosolic reducing power, which subsequently was used to drive the biochemical charging of the other energy carriers. By inspecting the flux-carrying reactions, we disrupted all EGCs by performing appropriate reaction deletions.

### Resolving the biosynthetic pathway of PUFAs

In thraustochytrids, the biosynthesis of PUFAs primarily occurs by the action of a PKS enzymatic complex. Here, the acyl chains are covalently attached to the acyl-carrier protein (ACP) domain of the enzyme complex, which directs the moiety to the various catalytic domains during successive rounds of chain elongation (Ratledge, 2004). The PKS pathway from the template model followed a more simplistic mechanism where the associated biosynthetic steps were merged into generic reactions involving erroneous metabolic intermediates. Additionally, the acyl moieties were present as acyl-CoA intermediates, thus omitting the final hydrolysis of the fatty acid from the ACP and subsequent condensation with CoA. We therefore implemented a more accurate representation of the PKS pathway in which the acyl-CoA intermediates were replaced by acyl-ACPs. We added every single catalytic step as unique reactions in order to depict the presumed biochemical reaction mechanism occurring *in vivo* (Ratledge, 2004). We also implemented the final hydrolysis of the acyl chain in order for the model to predict more realistic energy demands due to the ensuing ATP-driven condensation with CoA. The updated PKS pathway contained a total of 81 unique metabolic reactions, with 75 associated acyl-ACP intermediates. The biosynthetic pathway was able to synthesize all of the PUFAs that T66 accumulates during lipid accumulation (A N Jakobsen, Aasen, Josefsen, & Strom, 2008). These consist of the two *ω*-3 fatty acids EPA (c20:5(n-3)) and DHA (c22:6(n-3)), and the *ω*-6 fatty acid docosapentaenoic acid (DPA, c22:5(n-6)). The resulting pathway thus diverges early on in the biosynthetic process by either retaining the unsaturated *π*-bond at the n-3 position, generating EPA and DHA, or reducing it, eventually producing DPA.

In iCY1170_DHA, 15 separate genes were present in the associated gene rules of the PKS pathway. However, upon further inspection it was revealed that many of these enzymes contained rather generic PKS domains, which did not seem to constitute either of the three catalytic subunits of the PKS complex. Through the process of manual gene reannotation, we found three candidate genes in the genome of T66 which showed highly significant sequence similarity with the functionally characterized PUFA PKS subunits of *Thraustochytrium* sp. 26185 (Table 1) (Meesapyodsuk & Qiu, 2016). Along with an additional gene encoding the auxilliary phosphopantetheinyl transferase (acpT) subunit not present in iCY1170_DHA, we used these four genes to replace the previous set of 15 genes.

**Table 1:**
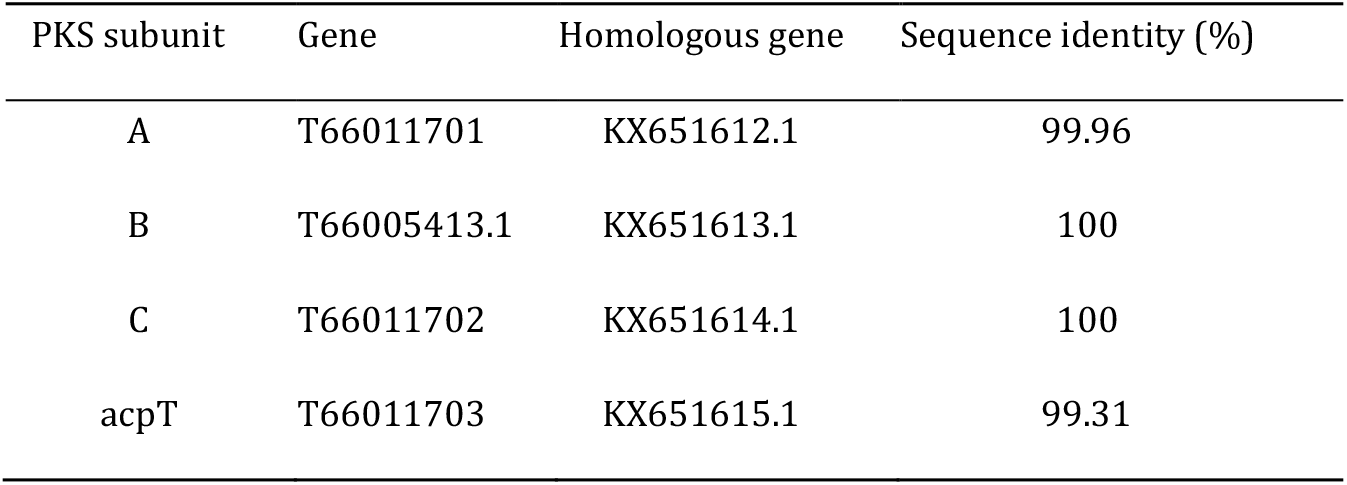
Candidate genes encoding the enzymatic subunits of the PKS complex responsible for the biosynthesis of PUFAs in T66. The homologous genes are GenBank accession numbers of functionally characterized proteins of *Thraustochytrium* sp. 26185 (Meesapyodsuk & Qiu, 2016).

### Addition of a peroxisomal compartment

Due to its central role in lipid metabolism, it is necessary to incorporate a peroxisomal compartment to achieve a high-fidelity reconstructed network. Following extensive literature reviews and identification of proteins that were hypothesized to be localized in the peroxisome, we added a peroxisomal compartment to the reconstruction. This compartment harbours 135 unique reactions, consisting of 92 internal and 43 transport reactions, respectively. Its metabolic capabilities are mainly associated with that of *β*-oxidation of fatty acids, along with the associated glyxoylate shunt, allowing for the gluconeogenic biosynthesis of carbohydrates from the acetyl-products of the *β*-oxidation pathway. The constituent reactions of the latter anaplerotic cycle were predicted to be localized to the peroxisome, as well as the cytosol and mitochondria, suggesting a rather complex interexchange of the metabolic intermediates between these compartments.

While the reactions associated with the glyoxylate shunt were present in iCY1170_DHA, thus allowing for growth on two-carbon compounds such as acetate, the model was unable to grow on any of its implemented fatty acids. This was caused by insufficient incorporation of necessary transport reactions and the erroneous subcellular localization of the mitochondrial electron-transferring-flavoprotein dehydrogenase (EC:1.5.5.1) to the cytosol, consequently preventing the mitochondrial degradation of these acyl chains. Additionally, indispensable enzymatic steps required for the oxidation of unsaturated fatty acids, such as Δ^3^-Δ^2^-enoyl-CoA isomerase (EC:5.3.3.8) and 2,4-dienoyl-CoA reductase (EC:1.3.1.34), were also not implemented, thus preventing the metabolic network from being able to degrade the collection of fatty acids that it is able to synthesize. When the required reactions were added along with the peroxisomal compartment, the model was able to grow on all implemented fatty acids, as well as a set of volatile fatty acids reported to support growth of T66 cultures (Table 2) (Patel, Rova, Christakopoulos, & Matsakas, 2020).

**Table 2:** Qualitative assessment of growth using the draft model, iVS1191, and iCY1170_DHA on various longto very-long chain fatty acids, as well as a set of volatile fatty acids (formate, acetate, propanoate, butyrate, pentanoate, and hexanoate) which has been shown to support growth of T66 *in vivo* (Patel et al., 2020). Uptake rates were arbitrarily set to 10 mmol gDW^-1^ h^-1^. Simulated *in silico* growth and non-growth is denoted by + and -, respectively.

| Carbon source | Draft model<sup>†</sup> | iVS1191 | iCY1170_DHA |
| --- | --- | --- | --- |
| Myristic acid, c14:0 | - | + | - |
| Palmitic acid, c16:0 | - | + | - |
| Palmitoleic acid, c16:1(n-7) | - | + | - |
| Stearic acid, c18:0 | + | + | - |
| cis-Vaccenic acid, c18:1(n-7) | - | + | - |
| EPA, c20:5(n-3) | - | + | - |
| DPA, c22:5(n-6) | - | + | - |
| DHA, c22:6(n-3) | - | + | - |
| Formate, c1:0 | - | + | - |
| Acetate, c2:0 | + | + | + |
| Propanoate, c3:0 | - | + | - |
| Butyrate, c4:0 | - | + | - |
| Pentanoate, c5:0 | - | + | - |
| Hexanoate, c6:0 | - | + | - |

<sup>†</sup>Draft model following the initial gap filling to enable biomass production.

Based on available genomic data, the *β*-oxidation of fatty acids in T66 is situated in both the peroxisomal and mitochondrial compartment. However, the chain-length specificities of these two enzymatic machineries remain to be elucidated. We therefore made the decision, similar to that of the GSM of *Phaeodactylum tricornutum* iLB1027 lipid (Levering et al., 2016), in which acyl chains of length 20 and more were assumed to initially be degraded solely in the peroxisomal compartment. The presumed end product octanoyl-CoA is then exported out of the peroxisome for further degradation in the mitochondrial *β*-oxidation pathway due to the presence of a peroxisomal carnitine O-octanoyltransferase (EC:2.3.1.137). The genuine substrate specificities of the two *β*-oxidation pathways are most likely overlapping. However, barring solid biochemical evidence of their acyl-chain preferences, we deem this differentiation the most probable based on the available genomic information.

### Cellular biomass reformulation

The biomass objective function is an abstract reaction in a GSM which contains the necessary biomolecular components to generate a unit of cellular dry weight (Feist & Palsson, 2010). By performing appropriate stoichiometric scaling of each biomass component, the flux through this reaction directly corresponds to the specific growth rate (h^-1^) of the organism (Chan, Cai, Wang, Simons-Senftle, & Maranas, 2017). Capturing the accurate macromolecular composition within the biomass reaction is therefore essential in order to correctly predict growth rates using a reconstructed GSM (Feist & Palsson, 2010). During the initial stages of the draft model construction and refinement, we used the biomass reaction of the template model iCY1170_DHA. However, the large content of lipids (43%, mainly TAGs) and carbohydrates (32%), and low content of proteins (12%), suggests a biomass deprived of nitrogenous compounds, presumably being closer to that of lipid accumulation rather than exponentially growing cells. In strain T66, the lipid content of exponentially growing cells is merely 13%, not reaching the above-mentioned levels until late in the lipid accumulation phase (A N Jakobsen et al., 2008). Additionally, the most prevalent or only lipid class found in exponentially growing thraustochytrids with excess nitrogen are polar lipids (Aasen et al., 2016).

Consequently, we found it prudent to reformulate the biomass composition reaction using parts of the macromolecular composition data from a recently published GSM of the thraustochytrid strain *Oblongichytrium* sp. RT2316-13 (Shene et al., 2020). This biomass reaction is specifically tailored for exponentially growing cells where the lipid proportion is free of neutral lipids such as TAG, and predominately contains the polar lipids phosphatidylcholine and phosphatidylethanolamine. Additionally, the protein content has a more biologically realistic value of ~49%. Using an approach similar to that of previous studies (Levering et al., 2016), we added biomass-precursor reactions, each containing sets of molecules belonging to one of the particular categories: proteins, carbohydrates, DNA, RNA, unsaponifiable matter, polar lipids, and free fatty acids. We implicitly added growth-associated maintenance (GAM) requirements by choosing alternative biomass precursor metabolites and by adding associated polymerization costs (e.g. nucleoside triphosphates for RNA synthesis). We also constrained the non-growth associated maintenance (NGAM) reaction to 9.9 mmol gDW^-1^ h^-1^, assuming a linear relationship between the cultivation temperature and maintenance requirements using model-fitted values previously published (Shene et al., 2020). The biomass composition and detailed calculations are available in ‘SuppInfo Biomass composition’.

### Final model reconstruction

The final model reconstruction iVS1191 contained a total of 1657 metabolites, 2095 reactions (1449 metabolic reactions, 415 transport reactions and 232 exchange reactions), and 1191 associated genes. Each of these reactions are associated with 82 unique subsystems, which are subdivided into five individual compartments, one external extracellular compartment, as well as four internal compartments; the cytoplasm, the peroxisome, the mitochondria, and the mitochondrial intermembrane space. One of the most prominent changes from the template model iCY1170_DHA is the additional number of reactions, where the number of transport reactions shows the largest increase. The modest increase in the number of genes can be explained by the process of manual curation. Here, 233 genes from the draft model with sub par hits against enzymes of known functionalities was removed during the process. At the same time, 270 genes were added, either to replace those that were discarded, or as associated genes of novel metabolic reactions not present in iCY1170_DHA. Consequently, although the difference might seem insignificant, considerable alterations have been performed so as to more accurately reflect genuine biological organization with these gene-reaction associations.

In addition to iCY1170_DHA, a revised GSM of *S. limacinum*, herein referred to as iCS1079, was recently published to simulate the growth and lipid phenotype of the closely related thraustochytrid strain *Oblongichytrium* sp. RT2316-13 using dynamic FBA (Shene et al., 2020). Here, the authors modified iCY1170_DHA by manually curating the model constituents and adding 220 new reactions to the reconstruction associated with amino acid, carbohydrate, and lipid metabolism. Using these previously reconstructed models, we compared their model components and properties with that of iVS1191. As presented in Table 3, iVS1191 shows a significant increase in metabolic coverage compared to both iCY1170_DHA and iCS1079. Specifically, we accomplished a considerable reduction in the number of blocked reactions and associated dead-end and orphan metabolites. In both iCY1170_DHA and iCS1079, a substantial subsection of the reactions were unable to carry a non-zero flux under any condition, constituting around 44% and 49% of all model reactions, respectively. Similarly, around 38% and 35% of the metabolites were categorized as metabolic orphans or dead-ends, either only being consumed or produced by the models. Even with a substantial increase in the number of reactions in iVS1191, only around 16% of the reactions were found to be blocked, while merely 16% of the metabolites were classified as metabolic orphans or dead-ends. This demonstrates how iVS1191 is able to utilize a greater subset of its metabolic capability, expectedly giving rise to more accurate phenotypic predictions.

**Table 3:** Comparison of features of the final model reconstruction iVS1191 and the two published thraustochytrid GSMs iCY1170_DHA and iCS1079. The blocked reactions were identified by having minimal and maximal fluxes of zero when running flux variability analysis (FVA) on an open model (i.e. all exchange reactions left unconstrained). Dead-end and orphan metabolites were identified as metabolites unable to either be consumed or produced by the model. The ORF coverage was calculated by finding the fraction of protein-encoding genes in the model to the total number of predicted ORFs of the organism.

| Features | iVS1191 | iCY1170_DHA | iCS1079 |
| --- | --- | --- | --- |
| Compartments | 5 | 3 | 4 |
| Genes | 1191 | 1170 | 1079 |
| Metabolites | 1657 | 1659 | 1454 |
| Reactions | 2095 | 1769 | 1610 |
| *Metabolic* | 1448 | 1385 | 1352 |
| *Transport* | 415 | 195 | 178 |
| *Exchange* | 232 | 189 | 80 |
| Blocked reactions | 332 | 775 | 789 |
| Dead-end/orphan metabolites | 265 | 628 | 512 |
| ORF coverage (%)<sup>†</sup> | 10.2 | 7.9 | 7.3 |
| MEMOTE score (%) | 90 | 28 | 39 |

<sup>†</sup>Genes found in the nucleotide genome sequence, but not as unique ORFs, were also included to account for insufficient coverage of the predicted protein sequences.

While the ORF coverage of iVS1191 did show a pronounced increase from that of iCY1170_DHA and iCS1079 (Table 3), this is most likely an effect of a smaller genome, and a corresponding reduction in the number of predicted ORFs. Whereas the genome size of *S. limacinum* is 60.93 Mb, with 14,859 unique ORFs (Institute, 2017), the genome size of T66 is merely 43 Mb, with a corresponding set of 11,683 predicted ORFs (Liu et al., 2016). Although this could imply a deficient genome coverage, it is comparable to other high-quality GSMs of the oleaginous species *Mortierella alpina* (Chao Ye et al., 2015) and *Yarrowia lipolytica* (Pan & Hua, 2012), each possessing an ORF coverage of 9.5% and 9.6%, respectively (C Ye et al., 2015).

We also benchmarked iVS1191 against the open-source software suit MEMOTE (Lieven et al., 2020), which provides a collection of consensus model tests that evaluate key properties such as stoichiometry, annotation, biomass, and formal correctness of the model. When compared with the other thraustohcytrid GSMs, iVS1191 showed major improvements in all test categories, obtaining a final score of 90% compared to 28% and 39% for iCY1170_DHA and iCS1079, respectively. Of particular importance was the considerable improvement in model consistency, which evaluates the extent of stoichiometric inconsistency, metabolic connectivity, and mass and charge balance. In this category, iVS1191 was awarded a score of 96%, while iCY1170_DHA and iCS1079 obtained a score of 64% and 87%, respectively.

### Phenotypic growth validation

A key part of the network evaluation stage is to compare model predictions with experimental data. Uncovered disparities might highlight deficiencies within the model reconstruction and further guide the modeler in subsequent reformulations and updates to close in on the gap between experimental and predicted phenotypes (Thiele & Palsson, 2010). To assess the quality of the constructed model, we first compared the predicted specific rate of biomass production against growth rates obtained from batch cultures of T66 on aerobic, minimal carbon-limited medium. We incorporated measured specific substrate-uptake rates as lower bounds on the exchange reactions of the corresponding model metabolites and optimized growth by maximizing the biomass objective function using the standard FBA formulation of Eqs. (1)-(3). As presented in Table 4, the model-simulated growth-phenotypes were found to be highly consistent with the growth rates determined experimentally on both glucose and glycerol. The predicted growth on glucose showed a relative error of merely −1.6%, while the relative error of the simulated growth on glycerol was only 3.5%.

**Table 4:** Comparison of *in vivo* and *in silico* specific growth rates (h^-1^) on aerobic carbon-limited minimal media, using measured carbon uptake fluxes (mmol gDW^-1^ h^-1^) to constrain the metabolic model. Growth rates were predicted by optimizing for biomass production.

|  | Minimal glucose | Minimal glycerol |
| --- | --- | --- |
| Carbon uptake *in vivo* | 1.440 | 2.440 |
| Growth *in vivo* | 0.124 | 0.114 |
| Growth *in silico* | 0.122 | 0.118 |
| Ammonium uptake *in vivo* | 1.110 | 0.990 |
| Ammonium uptake *in silico* | 0.981 | 0.953 |

## Discussion

With this work, we present a successfully reconstructed high-quality GSM of the thraustochytrid strain *Aurantiochytrium* sp. T66 termed iVS1191. By complementing the use of automatic tools for model reconstruction with extensive and comprehensive manual curation, we considerably expanded the metabolic scope and coverage from that of the template reconstruction iCY1170_DHA. In doing so, the connectivity of the metabolic network showed drastic improvements when compared to this previously reconstructed GSM, as well as the recently published thraustochytrid model iCS1079, significantly reducing both the number of blocked reactions and associated dead-end and orphan metabolites. Model-driven analysis of growth demonstrated that iVS1191 could grow on all the fatty acids that it accumulates (A N Jakobsen et al., 2008), as well as a set of volatile fatty acids shown to enable growth *in vivo* (Patel et al., 2020), which both iCY1170_DHA and the draft reconstruction failed to utilize as single carbon sources (Table 2). Understanding the role of this degradation of lipids and fatty acids during lipid accumulation is essential when trying to discern the metabolic properties of lipid production in thraustochytrids. The reconstructed model thus provides a highly valuable tool as a starting-point for model-driven generation of genetic engineering and cultivation strategies for increased DHA and lipid synthesis in T66.

During the model reconstruction process, we subjected the model to considerable manual curation and refinements. Particularly, we performed a genome-wide gene re-annotation of all the metabolic proteins of the two draft reconstruction. The motivation behind this labour-intensive re-annotation and comprehensive manual curation was threefold. Firstly, during the manual gap filling procedure, as well as throughout the identification of novel metabolic capabilities, the quality of iCY1170_DHA was questioned due to the apparent scarcity of manual curation of the model reactions and associated genes. Several genes seemed to be annotated to erroneous sets of reactions, and the subcellular localization of a large number of reactions seemed to be based purely on automatic predictions. A telling example was the subcellular localization of the electron-transferring-flavoprotein dehydrogenase (EC:1.5.5.1), which transfers electrons from multiple mitochondrial flavoprotein dehydrogenases, thus coupling the degradation of fatty acids and certain amino acids with oxidative phosphorylation (Watmough & Frerman, 2010). In iCY1170_DHA, this reaction was localized to the cytosol, effectively preventing the model from properly degrading these metabolites (see Table 2). Secondly, during the comparison between the genes of the initial draft model and the draft reconstruction from KEGG, considerable disparities in associated reactions were found between the genes that were present in both models. This was particularly true for proteins containing multiple catalytic domains, where the KEGG functionality in RAVEN appeared to have difficulties assigning appropriate KO IDs to these multi-functional enzymes. The quality of the reconciled draft models was therefore greatly assisted by this manual re-annotation and consolidation process. Lastly, inferring enzymatic activity from the genome annotation of T66 was in many cases rather challenging, primarily because of limited biochemical information in the annotation, but also due to occurrences of inaccurate assignments of database identifiers as well as associations with deprecated database entries.

Owing to this extensive curation of the individual model components, the stoichiometric consistency, metabolic connectivity, and mass and charge balance of the reconstruction exhibited drastic advancements. These curations allowed for the inclusion of a peroxisomal compartment, as well as a more realistic implementation of the PUFA-generating PKS pathway. Concurrent changes in the subcellular localization of key proteins associated with the *β-* oxidation of fatty acids enabled the model to grow on all its endogenously synthesized fatty acids, as well as multiple experimentally verified carbon sources (Table 2). Comparison of quantitative *in vivo* and *in silico* growth phenotypes also revealed the model able to accurately predict specific growth rates on both minimal glucose and glycerol media (Table 4), suggesting that the implemented biomass composition and associated ATP maintenance requirements are close to being representative of exponentially growing cells of T66. However, the predicted *in silico* ammonium uptake rates did show a larger deviance from the experimental values, particularly for growth on minimal glucose media in which the relative error was found to be −11.6% (Table 4). This indicates that the content of nitrogenous compounds of the biomass composition is somewhat insufficient, thus leading to a lower requirement for exogenously supplied ammonium. These results highlights the necessity for strain-specific and condition-dependent biomass compositions which in the future can be implemented with the reconstructed GSM to improve its predictive capabilities.

## Conclusion

We have developed and constructed a high-quality GSM of the thraustochytrid *Aurantiochytrium* sp. T66 termed iVS1191. The presented model provides the most comprehensive biochemical knowledgebase of any thraustochytrid to date, with a particular emphasis on lipid metabolism. Subjecting the model to large-scale manual curation, we significantly improved the metabolic coverage and scope from that of previously published thraustochytrid GSMs. With a reformulated biomass composition, the model was able to accurately predict specific growth rates on minimal carbon media, as well as qualitatively reproduce growth phenotypes on multiple experimentally tested nutrient sources. We strongly believe that the reconstructed model will form a solid framework to explore and understand the metabolic properties of strain T66 and aid in the *in silico* generation of metabolic engineering and cultivation strategies for improved lipid and *ω*-3-PUFA productivities. Additionally, the model encompasses a quality-controlled and quality-assured scaffold for future reconstructions of strain-specific thraustochytrid GSMs of great importance for biotechnological and industrial applications.

## Supporting information

SuppInfo_Biomass_Composition

SuppInfo_Dissipation_reactions

SuppInfo_Figure_1

SuppInfo_Figure_1

SuppInfo_Gene_reannotation

SuppInfo_Table_2

SuppInfo_Table_3

SuppInfo_Table_4

SuppInfo_draftModel

SuppInfo_iCS1079

SuppInfo_iCY1170_DHA

SuppInfo_iVS1191

SuppInfo_keggModel

## Acknowledgements

The authors would like to thank T. Kumelj and C. Schulz (Norwegian University of Science and Technology) for helpful discussions during the project, as well as S. Marci šauskas (Chalmers University of Technology) for assistance with RAVEN and the initial model reconstruction. We would also like to thank Inga Marie Aasen (SINTEF) for conducting the growth experiments of T66.

