## Supplementary material for "High-quality genome-scale metabolic model of *Aurantiochytrium* sp. T66": SuppInfo_Dissipation_reactions

Table S1: Set of dissipation reactions used to identify thermodynamically infeasible EGCs. Each dissipation reaction were iteratively added to the model, selected as the objective function, and subsequently maximized when all exchange reactions were constrained to zero. Any non-zero optimal objective value indicated the presence of an ECG.

| Energy carrier | Dissipation reaction |
|---|---|
| ATP | $ATP + H_2O \longrightarrow ADP + Phosphate + H^+$ |
| CTP | $CTP + H_2O \longrightarrow CDP + Phosphate + H^+$ |
| GTP | $GTP + H_2O \longrightarrow GDP + Phosphate + H^+$ |
| UTP | $UTP + H_2O \longrightarrow UDP + Phosphate + H^+$ |
| ITP | $ITP + H_2O \longrightarrow IDP + Phosphate + H^+$ |
| NADH | $NADH \longrightarrow NAD^+ + H^+$ |
| NADPH | $NADPH \longrightarrow NADP^+ + H^+$ |
| FADH_2 | $FADH_2 \longrightarrow FAD + 2\,H^+$ |
| Ubiquinol-9 | $Ubiquinol-9 \longrightarrow Ubiquinone-9 + 2\,H^+$ |
| Acetyl-CoA | $Acetyl-CoA + H_2O \longrightarrow acetate + CoA + H^+$ |
| L-glutamate | $L-glutamate + H_2O \longrightarrow 2\text{-oxoglutarate} + Ammonium + 2\,H^+$ |
