## Supporting Information Legends for "High-quality genome-scale metabolic model of *Aurantiochytrium* sp. T66"

**SuppInfo_Biomass_Composition.xlsx**

Detailed calculations of the biomass composition used in iVS1191.

**SuppInfo_Dissipation_reactions.pdf**

Set of dissipation reactions used to identify the thermodynamically infeasible energy-generating cycles (EGCs).

**SuppInfo_Gene_reannotation.xlsx**

List of manual gene re-annotation of all the included metabolic genes in iVS1191.

**SuppInfo_Figure_1.txt**

MATLAB script used to reproduce Figure 1 in the main text of the article.

**SuppInfo_Figure_1.xlsx**

Corresponding data used to reproduce Figure 1.

**SuppInfo_iCS1079.xml**

Model file of iCS1079 in xml format.

**SuppInfo_iCY1170_DHA.xml**

Model file of iCY1170_DHA in xml format.

**SuppInfo_iVS1191.xml**

Model file of iVS1191 in xml format.

**SuppInfo_draftModel.xml**

Model file of the T66 draft model in xml format.

**SuppInfo_keggModel.xml**

Model file of the T66 KEGG model in xml format.

**SuppInfo_Table_2.txt**

MATLAB script used to reproduce Table 2 in the main text of the article.

**SuppInfo_Table_3.txt**

MATLAB script used to reproduce Table 3 in the main text of the article.

**SuppInfo_Table_4.txt**

MATLAB script used to reproduce Table 4 in the main text of the article.
